# Paired evaluation defines performance landscapes for machine learning models

**DOI:** 10.1101/2022.09.07.507020

**Authors:** Maulik K. Nariya, Caitlin E. Mills, Peter K. Sorger, Artem Sokolov

**Author notes:** Corresponding author cc.

## Abstract

The true accuracy of a machine learning model is a population-level statistic that cannot be observed directly. In practice, predictor performance is estimated against one or more test datasets, and the accuracy of this estimate strongly depends on how well the test sets represent all possible unseen datasets. Here we present paired evaluation, a simple approach for increasing the robustness of performance evaluation by systematic pairing of test samples, and use it to evaluate predictors of drug response in breast cancer cell lines and of disease severity in patients with Alzheimer’s Disease. Our results demonstrate that the choice of test data can cause estimates of performance to vary by as much as 30%, and that paired evaluation makes it possible to identify outliers, improve the accuracy of performance estimates in the presence of known confounders, and assign statistical significance when comparing machine learning models.

## INTRODUCTION

Effectively evaluating the performance of predictive computational models is a crucial aspect of machine learning. Knowing when a model is accurate allows for reliable predictions on new data and provides valuable insights about which features in the training data carry predictive information. However, the true accuracy of a model is a population level statistic that is generally unknown, because it is impossible to consider all — potentially infinitely many — datasets to which a model will be applied. Model performance must therefore be estimated by appropriately sampling available data, and reliable estimates require a sufficient number of points to adequately represent the population. The presence of systematic biases and confounding variables can lead to incorrect accuracy estimates and inflated confidence in machine learning models that are subsequently found to perform poorly in deployment^1^. This is closely related to the well-known issue of overfitting^2^, whereby a model trained on one set of data points fails to generalize to a new set of data. Conventional methods for performance evaluation can fail to detect overfitting when the same biases are present in training and test data. Robust performance estimates must therefore detect and account for these biases to accurately represent how the model would behave in the larger space of all possible data points.

When data are limited (as they commonly are in biomedicine), model performance is routinely evaluated using cross-validation, which involves withholding a portion of the available data and using the remainder to train a model, which is then evaluated against the withheld portion^3,4^. A widely used variant is *k*-fold cross-validation, which partitions available data into *k* equivalent fragments and rotates which fragment is withheld during training and used for evaluation. A specific and common example of this is leave-one-out cross-validation (LOOCV), where *k* is set to the number of data points. Other common approaches to cross-validation include Monte Carlo methods, in which a fixed proportion of data are repeatedly sampled and withheld for evaluation; and bootstrap methods, in which sample data with replacement are used to generate a training set, thus automatically defining the withheld partition based on the data points not sampled^5^. In their standard formulations, none of these methods explicitly account for the presence of systematic biases and confounders in the data, and model accuracy estimates obtained by these methods may not always reflect true predictor performance, particularly when dataset sizes are small.

The limitation of data availability is particularly prominent in -omics datasets, which commonly contain many molecular measurements (ca. 10^4^ for genome-scale data) from a relatively small number of samples (10-100). While conducting more experiments to increase sample size is sometimes possible, the small-sample issue is insurmountable in other cases due to the limited availability of biological material (number of available patient specimens for example) and the significant cost associated with molecular profiling. For example, cell culture studies focused on breast cancer are generally limited to the ∼75 commercially available breast cancer cell lines. While deriving new cell lines is possible, it is time consuming and expensive^6^. Moreover, new lines potentially suffer from the same limitations as existing lines with respect to confounders. The discrepancy between the low number of samples and the large number of molecular features available for any one sample introduces *low signal* scenarios. For example, the availability of deep gene expression data that cover thousands of expressed genes in a small number of samples makes it difficult to detect relevant transcriptional changes in the overall expression variance^7^. A low number of samples can also lead to stratification bias, because it is not always possible to partition a small but discrete number of data instances into cross-validation folds in a way that preserves the statistical properties of the entire dataset in each fold^8^. Together, these issues represent a substantial challenge in making accurate estimates of performance for models trained on - omics and similar datasets.

In addition to challenges arising when sample number is low, biological and clinical datasets often contain both known and unknown confounding relationships among variables. For example, a recent study found that the dominant signal in a prototypical large multi-center drug-response screen aligned with the location at which the data was collected and not the drug or cell line^9^. Knowing when a machine learning model inadvertently learns to recognize such a lurking variable can help prevent spurious correlations and erroneous conclusions. A popular approach for dealing with confounding and lurking variables is to modify the input data in a way that removes or reduces their effect, as implemented by ComBat^10^, Surrogate Variable Analysis^7^, Removal of Unwanted Variation^11^, and linear models for microarray data^12^. However, modification of the original data can inadvertently introduce new artifacts that erroneously amplify differences between data groups and inflate estimates of model performance^13^. Some batch-correcting methods also assume an underlying statistical distribution for the data, making them inappropriate for scenarios in which the data distribution is unknown.

In this work, we describe *paired evaluation*, a simple approach to model evaluation that systematically examines how a predictive model scores pairs of test data samples to generate a detailed decomposition of performance estimates. We demonstrate that paired evaluation can identify potential data outliers, assign statistical significance when comparing machine learning methods, and serve as a non-parametric method to accurately estimate model performance in the presence of known confounders without requiring modification of the underlying data. We consider two prediction tasks that leverage real-world datasets with known confounders: prediction of drug sensitivity in breast cancer cell lines, which is confounded by subtype (clinical: hormone receptor positive, HER2 amplified, triple negative; and molecular: luminal, basal); and prediction of Alzheimer’s Disease (AD) severity in post-mortem brain specimens, which is confounded by an individual’s chronological age. We show that minor variations in how the test data are paired for evaluation can reveal significant effects hidden by traditional approaches to model evaluation, and that the exclusion of outliers detected by paired evaluation can impact model interpretation. Lastly, we address the potentially prohibitive quadratic complexity of paired evaluation by proposing formulations that scale linearly with the number of test data points.

## RESULTS

Throughout this work, we quantify model accuracy using a popular metric, the area under the receiver operating characteristic curve (AUC)^14^. In binary classification, the AUC can be interpreted as the probability that a randomly chosen positive sample is correctly ranked above a randomly chosen negative sample^15^. Motivated by this interpretation, we developed paired evaluation, an approach in which a test dataset is broken up into pairs of samples, and a predictor is evaluated based on whether it ranks each pair correctly. We define a pair of samples to be rankable if their labels can be ordered – given experimental error and other uncertainty – by the corresponding data representation, e.g., the temporal arrangement of events (disease progression or death) in a survival dataset. The fraction of pairs ranked correctly is a direct estimate of AUC (**Figure 1a**). Paired evaluation is agnostic to the underlying machine learning method and can be applied in any cross-validation setting that withholds at least two data points in each fold. A special case of this is leave-pair-out cross-validation (LPOCV), in which a separate model is trained for each test pair^16^. LPOCV is particularly relevant for small-sample datasets with low signal-to-noise ratios, because it has been shown to be less susceptible to stratification bias than other popular cross-validation schemes^15–17^.

**Figure 1:**
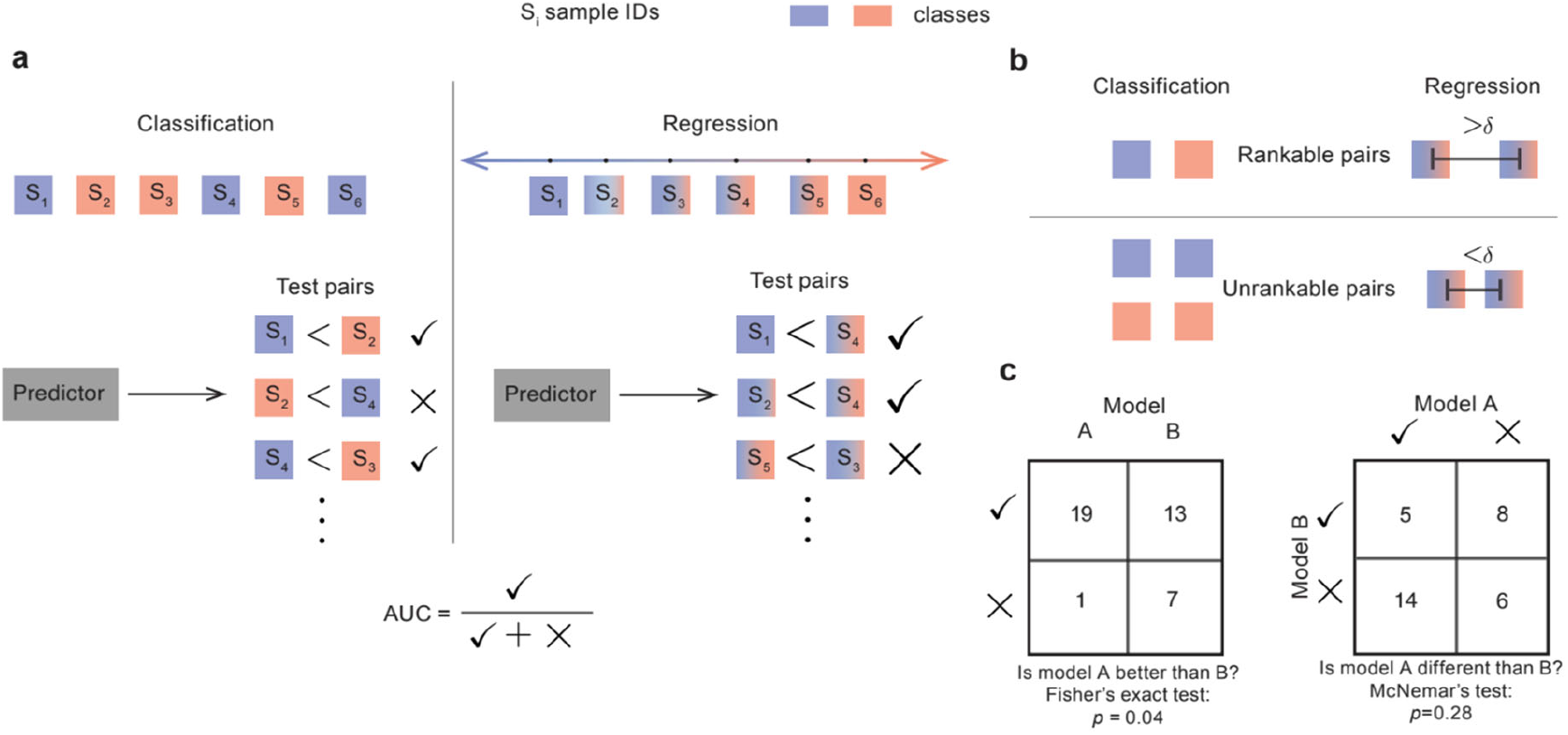
A schematic representation of paired evaluation. **a**. Individual samples in a test dataset are represented by squares, colored according to their true labels in binary classification and linear regression settings. The test dataset is broken up into rankable pairs, and a predictor is asked to score each pair separately. The scores are used to determine whether a given pair was ranked correctly (?) or incorrectly (X), and the AUC is determined by the fraction of correctly ranked pairs. **b**. The criteria for a valid rankable test pair. In binary classification, two samples are considered rankable if they belong to the opposite classes; in linear regression, a rankable pair of samples requires that the difference between their labels is greater than a predefined meta-parameter *δ*. **c**. An example comparison of two models, A and B. A 2 × 2 contingency table tallies the correctly and incorrectly ranked pairs by each model. Statistical significance of the difference in method performance is assessed by Fisher’s exact test and McNemar’s test.

Paired evaluation is not limited to binary classification and can be applied to any machine learning task that allows for an ordering of sample labels, including linear and ordinal regression, recommender systems, and survival analysis. To account for instrument error and other sources of uncertainty in -omics datasets, we introduce an optional meta-parameter *δ* that sets a minimum required distance of separation in the label space for a pair of samples to be considered rankable (**Figure 1b**). In all settings, AUC is estimated as the fraction of rankable pairs that are ranked correctly by a model. This estimate can be viewed as the average number of successes in a series of Bernoulli trials. While the trials are not independent and identically distributed (i.i.d.), the resulting AUC values will often follow a binomial distribution in practice and, based on the Central Limit Theorem, can be reasonably modeled with a Gaussian distribution when the number of pairs is sufficiently large.

The primary advantage of paired evaluation is that it provides a more detailed landscape of model performance than an AUC value alone since it allows models A and B to be compared against each other based on their ability to correctly rank individual pairs of datapoints. Specifically, paired evaluation enables assessment of statistical significance by simple construction of a two-by-two contingency table and utilization of the Fisher’s exact test (**Figure 1c**). Similarly, use of McNemar’s test makes it possible to detect instances in which two models perform differently even when their AUC values are comparable, which often signals that the models are complementary and suggests that aggregating their output with an ensemble model may lead to improved accuracy^18^.

### The choice of test data has a profound effect on estimates of model performance

Breast cancer is a heterogeneous disease that is clinically subtyped based on the levels of expression of three receptors: tumors expressing estrogen and/or progesterone receptors are classified as hormone-receptor positive (HR+), tumors overexpressing and/or amplified for the HER2 receptor tyrosine kinase are classified as HER2-positive, and those lacking expression of these three genes are classified as triple negative (TNBC). In practice, clinical subtype determines how a cancer will be treated. Breast cancers are also classified based on gene expression profiles into four intrinsic molecular subtypes: luminal A, luminal B, basal and HER2-enriched^19,20^. Molecular and clinical subtypes overlap but are non-identical. Given the high concordance between clinical and molecular subtypes in our cell line data (**Figure S5**), we followed the common practice of separating lines into luminal (HR+, HER2+) and basal (TNBC) molecular subtypes as a potential confounding variable ^21–24^.

We considered a dataset recently collected in our laboratory that characterizes the sensitivity of 63 breast cancer cell lines of different subtypes to 72 small molecule drugs, with a focus on kinase inhibitors. The dataset comprises growth rate-corrected measures of drug sensitivity (GR values^25^), determined using a microscopy-based assay of cell proliferation and death^26^, and pre-treatment transcriptional and proteomic^27^ profiles for each cell line. To demonstrate the effectiveness of paired evaluation, we considered a simple machine learning setup, in which random forest regression models were trained to predict drug sensitivity — measured as area over the growth rate curve (GR_AOC_) — from the baseline mRNA expression of a set of pre-selected genes (**FigureS1**).

To maximally utilize the small number of data points (cell lines), we applied paired evaluation using LPOCV, where a separate model was trained for each test pair. To account for possible measurement error, we considered a pair of cell lines to be rankable if the difference in the corresponding GR_AOC_ labels was greater than a specified value of the meta-parameter *δ* (**Figure 1b**). We observed that the value of this meta-parameter had a dramatic impact on the estimate of model performance, with some estimates varying by as much as 30% (**Figure 2a**). This finding reinforces the importance of choosing a test set that accurately represents potential future data that would be encountered by the predictor. Here, a large value of *δ* presents an “easy” prediction task, in which it is necessary only to distinguish cell line pairs having large differences in drug sensitivity. Such scenarios produce higher perceived model performance, but these estimates are artificially inflated relative to observed differences between cell lines in general and may not represent the true accuracy of the model.

**Figure 2:**
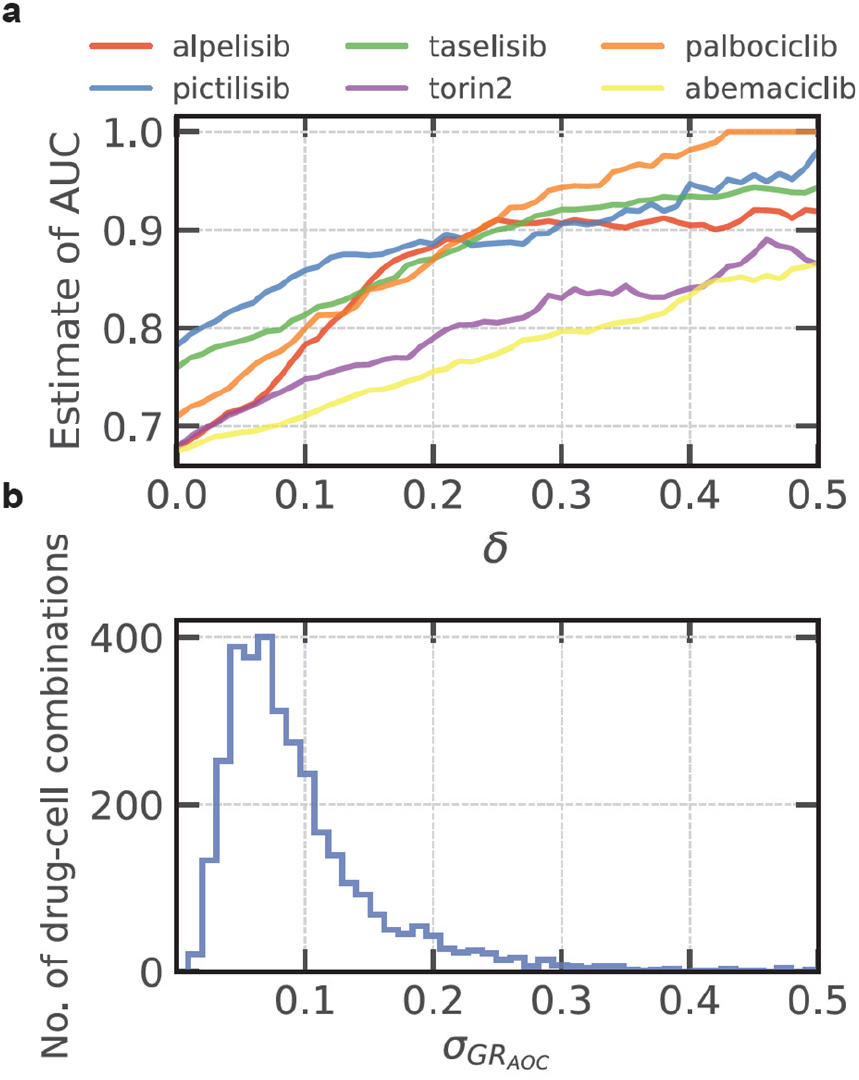
The composition of the set of rankable pairs plays a crucial role in evaluating predictive models of drug response in breast cancer cell lines. **a**. The parameter *δ* defines the set of rankable cell line pairs, which are then used to estimate AUC of random forest models in LPOCV. Each model was trained to predict drug sensitivity from baseline mRNA expression. Shown are estimates of performance for six select compounds **b**. The distribution of standard deviation in GR_AOC_ across technical triplicates for 3,400 drug-cell combinations. Predictive models are not expected to be able to distinguish two cell lines with GR_AOC_ values that lie within the corresponding standard deviation since it represents measurement error.

To establish a reasonable value for *δ*, we required that a corresponding model correctly ranks pairs of cell lines for which separation of GR_AOC_ values (labels) was greater than experimental error. For a given drug-cell line combination, the experimental error was taken to be the standard deviation of GR_AOC_ across three or four technical replicates. For any pair of cell lines, the larger of the two standard deviations was then used as the value for *δ* to determine if that pair was rankable. For most rankable pairs, this corresponded to a difference in GR_AOC_ of *δ* < 0.3 (the full range of GR_AOC_ values in our data was -0.7 to 1.9), with the total number of rankable pairs on the order of hundreds for each drug (**Figure 2b, Suppl. Table 1**).

### Effect of breast cancer subtype on model performance

In our dataset, the dominant variance in gene expression data and drug sensitivity for multiple drugs was observed to align with molecular subtype (**Figure S5**), consistent with previous studies^21,28^. Thus, subtype represents a known complication in the analysis of breast cancer drug response, and we sought to evaluate its impact on estimates of model performance. To accomplish this, we broadly classified cells as either luminal or basal and compared AUC estimates computed with all rankable pairs against estimates derived using only those rankable pairs for which both cell lines were of the same subtype. For many drugs, we observed a decrease in estimated AUC when the evaluation was performed on subtype-matched pairs (**Figure 3a**), suggesting that the corresponding predictors had learned to recognize molecular subtype as a confounder. Next, we estimated the correlation between drug sensitivity and subtype using one-way ANOVA and observed that the resulting F-statistic was a good indicator of the difference between AUC estimates (**Figure 3b**). Our results confirm that learned models place more emphasis on the molecular subtype when it is indeed a good predictive feature of drug sensitivity. However, when prediction is limited to a single subtype, the models are frequently less accurate. The balance between accuracy across subtypes vs. within a subtype must therefore consider the way in which a model will be used. For example, if a drug is approved only for one subtype, then a subtype-specific model may be what is required.

**Figure 3:**
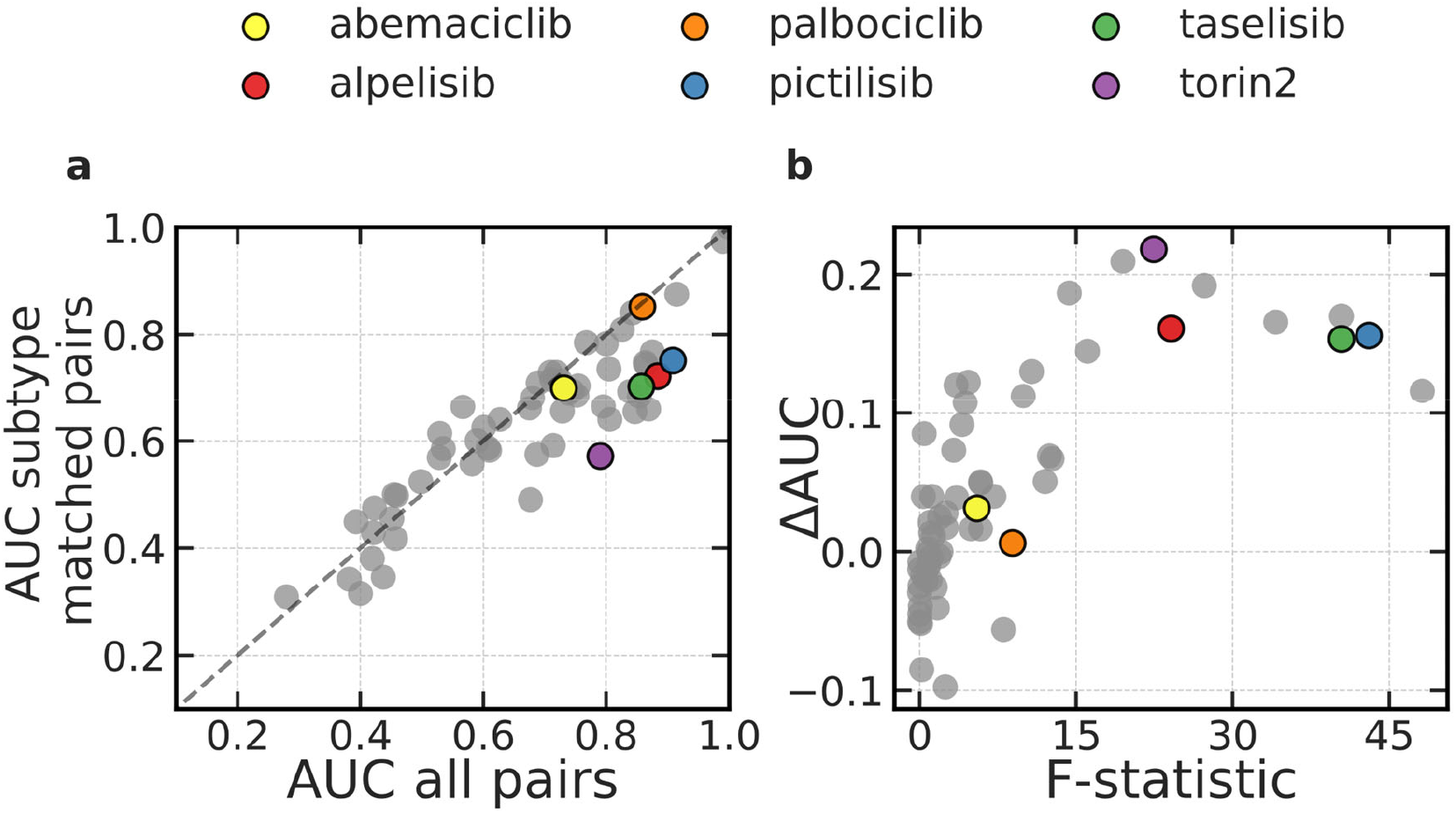
Effect of breast cancer subtype on the estimates of prediction accuracy. **a**. AUC estimates calculated using subtype-matched (y axis) and all (x axis) rankable pairs. The dashed line represents all hypothetical scenarios where the two AUC estimates agree. Each dot corresponds to one of 72 drugs. A subset of drugs is highlighted for closer examination. **b**. The difference between AUC estimates in panel b., computed as ΔAUC = AUC_all_ − AUC_subtype_ and plotted against matching one-way ANOVA to contrast GR_AOC_ distributions across breast cancer subtypes (as in panel a.). Each dot corresponds to one of 72 drugs.

To get a better understanding of the effect breast cancer subtype has on performance estimation, we considered six clinically relevant breast cancer drugs for closer examination (**Table 1**). Of these, alpelisib is currently approved for the treatment of HR+/HER2- metastatic breast cancers^29^ and is in clinical trials for HER2+ patients. Palbociclib and abemaciclib are approved for use in the same metastatic HR+ breast cancers with current attempts to expand the indication to TNBC and HER2^+^ disease^30^. Consistent with these clinical indications, we found that basal and luminal cell lines responded differently to alpelisib, pictilisib, taselisib, and Torin2, while no significant difference in response was observed for palbociclib and abemaciclib (**Figure S7**).

**Table 1:** Effect of breast cancer subtype on model performance.

| Drug |  | All rankable pairs | Subtype matched pairs | *p*-value |
| --- | --- | --- | --- | --- |
| alpelisib | ✓ | 337 | 80 | $8.81 \times 10^{-5}$ |
|  | ✗ | 30 | 24 |  |
|  | AUC | 0.92 | 0.77 |  |
| pictilisib | ✓ | 315 | 66 | $2.89 \times 10^{-4}$ |
|  | ✗ | 43 | 26 |  |
|  | AUC | 0.88 | 0.72 |  |
| taselisib | ✓ | 604 | 192 | $8.99 \times 10^{-9}$ |
|  | ✗ | 110 | 91 |  |
|  | AUC | 0.85 | 0.68 |  |
| torin2 | ✓ | 273 | 68 | $6.63 \times 10^{-8}$ |
|  | ✗ | 116 | 84 |  |
|  | AUC | 0.70 | 0.45 |  |
| palbociclib | ✓ | 367 | 176 | 0.90 |
|  | ✗ | 61 | 30 |  |
|  | AUC | 0.86 | 0.85 |  |
| abemaciclib | ✓ | 382 | 187 | 0.75 |
|  | ✗ | 177 | 82 |  |
|  | AUC | 0.68 | 0.70 |  |

In paired evaluation, the estimate of AUC was substantially lower for subtype-matched pairs when predicting sensitivity to alpelisib, pictilisib, taselisib, and Torin2 (**Table 1, Figure S6a**), suggesting that the corresponding predictors had at least partially learned to recognize the molecular subtype. In contrast, no statistically significant difference was observed when comparing AUC estimates made using all pairs and subtype-matched pairs for palbociclib or abemaciclib (**Table 1, Figure S6b**), two drugs whose sensitivity was more weakly correlated with subtype (**Figure S7**). Taken together, these findings suggest that when drug sensitivity is correlated with subtype, predictors implicitly learn features of the underlying subtypes. This may represent a desirable property in a setting where molecular subtype closely informs drug response.

### Detection and removal of outliers impacts model interpretation

In the current setting, model interpretation primarily involves inspecting feature importance scores to pinpoint genes that play a crucial role in determining drug response and resistance. Since the presence of outliers in the training data can skew feature importance scores, we investigated the effects of outlier removal on model interpretation. We asked if any rankable pair was more likely to be ranked incorrectly by a model if it included specific data samples. We were specifically concerned about outliers that arose from measurement error, or that were biologically very different from the norm. When predicting the sensitivity of breast cancer cell lines to Torin2, a polyselective mTOR/PIKK inhibitor^31^, we found that 526 out of 673 rankable pairs were ranked correctly by a random forest model (AUC estimate = 0.78). However, pairs containing the ZR7530 cell line were consistently ranked incorrectly (**Figure 4a,b**), suggesting that the cell line is an outlier. ZR7530 is a luminal cell line, and its gene expression profile clusters with profiles of other luminal cells (**Figure S2**). However, the cell line was more resistant to Torin2 than other luminal cell lines with a *GR*_*AOC*_ value more similar to that of TNBC lines, explaining the observed misranking of pairs containing ZR7530.

**Figure 4:**
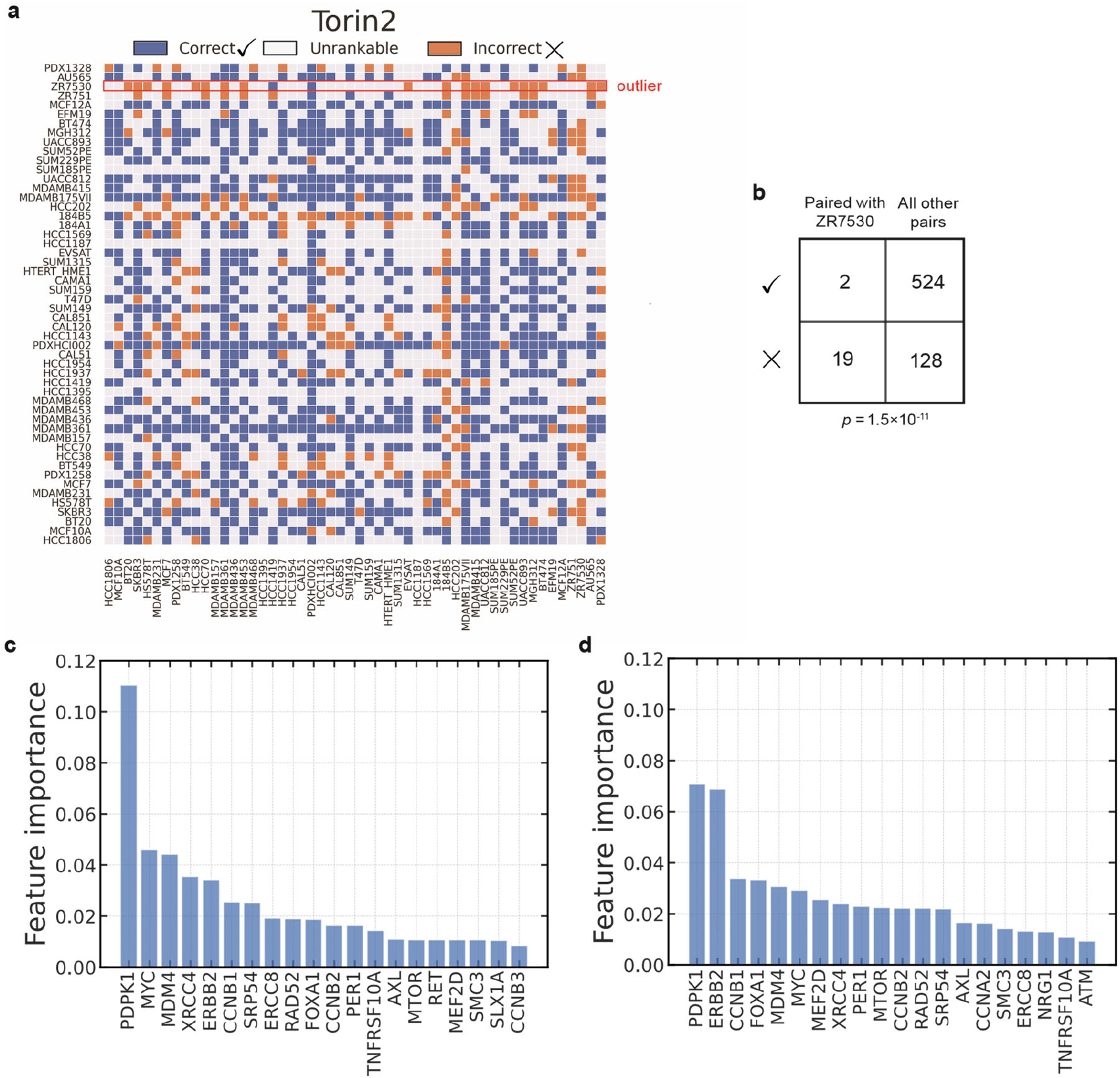
Paired evaluation detects outlier cell lines in the context of sensitivity to torin2. **a**. A performance landscape over all possible pairs of cell lines. A pair is colored blue if it was correctly ranked by the predictor and orange otherwise. Pairs that were not considered rankable because the corresponding *GR*_*AOC*_ values were not separated by the *δ* threshold are shown in gray. **b**. A 2×2 contingency table tallying correctly and incorrectly ranked pairs with and without the cell line ZR7530. The corresponding *p* value was computed with a Fisher’s exact test. **c**. Feature importance scores associated with a predictor trained on all cell lines. Shown are the top 20 features. **d**. Feature importance scores computed after removing the outlier ZR7530 and retraining the predictor on the remaining cell lines.

Removing ZR7530 from the dataset reduced the total number of rankable pairs to 652, of which 524 were ranked correctly (**Figure 4b**), leading to a small improvement in estimated model accuracy (AUC estimate = 524 / 652 = 0.8). When we compared feature importance scores before and after the removal of ZR7530, we observed that *ERRB2 (HER2)*, increased in importance (**Figure 4c,d)**, reconfirming that receptor status is heavily correlated with Torin2 response (**Table 1**). Similarly, we found that *MTOR*, which encodes a known target of Torin2^32^, also gains importance. More generally, these findings show that the feature importance of genes know to play an important role in breast cancer biology change when outliers are detected and removed in paired evaluation, and at least in some cases this increases interpretability.

We repeated the outlier analysis for all other drugs in our dataset and identified two other cases, corresponding to drugs E17 and palbociclib, for which sensitivity predictors consistently mis-ranked pairs containing a specific cell line. In both cases, removing the outlier led to a higher estimate of AUC, but the effect on feature importance varied. In the case of E17, the removal of outlier cell line MGH312 led to a substantial drop in the importance *CDKN2C* (**Figure S3**). However, removing the outlier cell line HCC202 from a predictor of palbociclib sensitivity did not have any substantial impact on feature importance (**Figures S4**). These results demonstrate that the presence of outliers in a dataset can lead to a consistently incorrect ranking of pairs, and the removal of these outlier can increase, decrease, or have no effect on feature importance, making paired evaluation a useful tool for model interpretation.

### Disease severity in Alzheimer’s decedents

Alzheimer’s Disease (AD) is a chronic neurodegenerative disorder that leads to memory loss and dementia. The disease is characterized by extracellular aggregates of the *β*−amyloid peptide and intracellular accumulation of hyperphosphorylated tau leading to neurofibrillary tangles (NFT). Several recent studies – those from the Accelerating Medicines Partnership - Alzheimer’s Disease (AMP-AD) program for example^33^ – have attempted to obtain molecular insight into disease mechanism using diverse omic datasets obtained from patient specimens; these data include whole genome sequencing, DNA methylation, mRNA and protein expression, and detailed clinical annotation.

Here, we consider the task of predicting disease severity from mRNA expression. We make use of the data collected by two joint longitudinal cohort studies, the Religious Orders Study (ROS) and the Memory and Aging Project (MAP), that comprise over two hundred bulk RNAseq profiles of postmortem brain specimens, along with matching pathology annotations^34,35^. We group data points into three categories based on Braak staging^36,37^: mild (Braak 1–2), moderate (Braak 3–4), and severe (Braak 5–6).

AD progression takes place on a timescale of years, and disease severity is strongly correlated with a patient’s age of death (AOD) (**Figure 5a)**. An important question is whether a predictor trained to recognize disease severity has instead learned to predict age, a situation that can lead to an overinflated estimate of performance and affect the interpretation of the genes and weights that make up the model. To address this, we used paired evaluation as a non-parametric way of evaluating the effect of a known confounder on performance estimates; this involved contrasting confounder-matched and confounder- mismatched rankable pairs. As with breast cancer data above, we used paired evaluation in a LPOCV setting, training a separate model for each rankable pair to make the most use of the limited amount of data.

**Figure 5:**
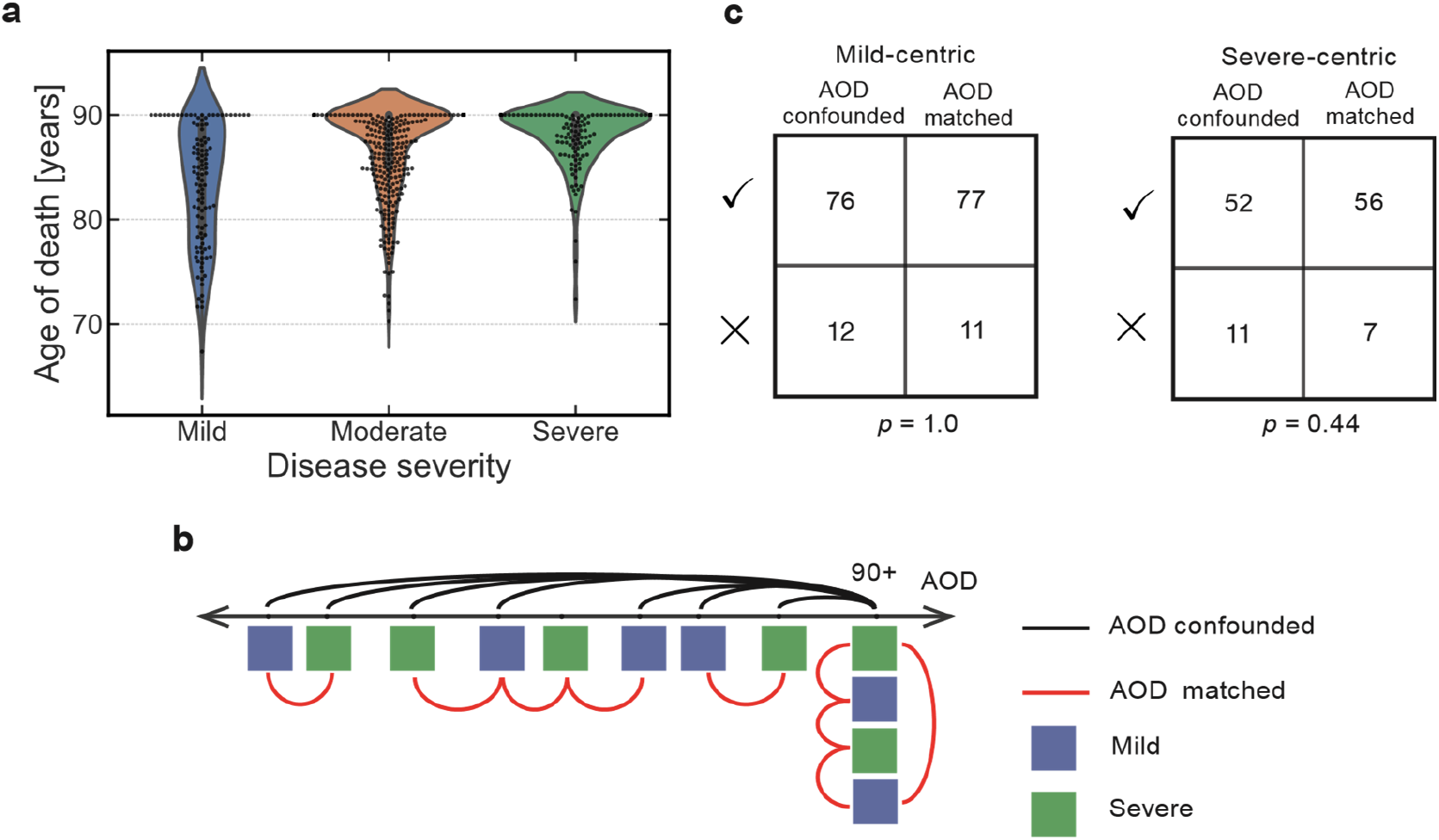
**a** The distribution of age of death (AOD) for patients who were diagnosed with mild, moderate or severe AD during postmortem pathology analysis. **b** Schematic representation of rankable pairs, selected to be either confounder-matched (red) or -mismatched (black). Each patient is represented by a square, colored according to the corresponding pathology annotation. The value of AOD is censored at 90 years of age in the dataset. **c** 2×2 contingency tables showing the correctly and incorrectly ranked test pairs for AOD confounded and AOD matched scenarios.

In our breast cancer analysis, the labels were continuous (varying from 0-1 GR_AOC_), and the confounder was represented by a discrete variable (basal or luminal subtype). The opposite is true of ROSMAP data; the labels are discrete, resulting in a straightforward definition of rankable pairs: two brain specimens are rankable, if they have distinct Braak stage ranges. Conversely, age is a continuous variable that is censored at 90 years old (y.o.). The censoring provides a natural inflection point for determining whether two data points are matched in age, giving rise to two evaluation scenarios (**Figure 5b**). In a scenario focused on mild disease (a mild-centric scenario), each individual who passed away before the age of 90 with mild disease was paired with individuals who had severe disease and was either closest in age (AOD-matched) or was chosen at random from the censored category of 90+ y.o. (AOD-confounded). Similarly, a severe-centric scenario pairs each 90+ y.o. patient who passed away with severe AD with a patient who had mild disease and was either randomly selected from the same 90+ y.o. category (AOD-matched) or the youngest patient in the cohort (AOD-mismatched). As a reference point, we also considered all possible rankable pairs.

In both mild-centric and severe-centric scenarios, each data point was associated with two rankable pairs that represent the minimal and maximal separation along the confounding variable (**Figure 5b)**. We trained logistic regression models to recognize disease severity from the corresponding mRNA expression profiles and applied paired evaluation to estimate model performance with each set of rankable pairs. We found that the models performed similarly for AOD-matched and AOD-confounded pairs (**Figure 5c**), and that both performance estimates were consistent with the one derived on all rankable pairs (mild- centric AUC = 0.87, severe-centric AUC = 0.85). Similar performance trends were also observed for the mild-vs-moderate and moderate-vs-severe classification tasks.

The analysis reveals that the presence of confounders does not necessitate that a predictor will learn to recognize them instead of the variable of interest. Paired evaluation provides a simple way to detect whether such situations occur and can facilitate decisions about when it is necessary to correct for confounding variables.

## DISCUSSION

We present *paired evaluation*, a method for deriving detailed landscapes of predictor performance for machine learned models based on the concept of rankable pairs of datapoints. We show how systematic pairing of data points can account for known confounders and identify outliers. We also show that statistical significance of model comparison can be assessed using contingency tables. While we made use of LPOCV in the current work due to the limited number of samples in our datasets, paired evaluation can be applied in any cross-validation setting that withholds at least two samples for testing in each fold.

The choice of test data can have a dramatic effect on estimates of model performance^1^. To get an accurate performance estimate, a test set must be a faithful representation of future data that a predictor might encounter in deployment. We therefore recommend that rankable pairs be defined using experimental knowledge and domain expertise. For example, in regression problems, the choice of a minimal difference (in a continuous variable) for a pair to be rankable (*δ*) could be based on variance across biological or technical replicates (**Figure 1b**); two data points that fall within this variance are routinely indistinguishable. Confounding and lurking variables are ubiquitous, but their presence may not be a drawback if they are biologically relevant and can assist in model interpretation. In the case of breast cancer cell lines, the difference between HR^+^, HER2^+^ and TNBC status confounds modeling of drug sensitivity but is informative for drugs that inhibit HR and HER2^28^. Conversely, a predictor that unintentionally learns to recognize what institution a subset of data was collected at in a multi-center study^9^ is unlikely to produce meaningful biological insight. Because confounders can have either positive or adverse effects on model interpretation, it is imperative to know when predictors have learned to recognize confounders. We show that paired evaluation is an effective, non-parametric method to detect this through simple comparison of performance values computed on confounder-matched vs. confounder-mismatched pairs. Importantly, paired evaluation achieves this without modifying the original data.

Our study has several limitations. In its present formulation, defining confounder-matched rankable pairs requires that the confounder values are known; however, many datasets may have unknown or unmeasured lurking variables that introduce unwanted batch effects^7^. To evaluate the effect of these hidden variables on the estimate of performance using paired evaluation, a user would first have to detect them using an external method. We also expect that information about hidden batch effects may be encoded in pairwise comparison of data points. Our future work will extend the outlier detection to identify groups of samples that exhibit similar misranking patterns as a method for approximating shared unobserved characteristics. While paired evaluation can detect situations in which confounders affected model training, the method provides no intrinsic means to correct for this effect, since the original training data is not modified. Furthermore, it is not trivial to delineate what aspect of model interpretation (*e*.*g*., feature importance) aligns with a confounder versus the variable of interest, even when paired evaluation signals that a predictor learned to recognize that confounder. Thus, paired evaluation represents the initial step in identifying potentially problematic confounder variables and outlier samples, but resolution of these may require other methods.

## METHODS

### Estimation of AUC

We consider a pair of samples *i* and *j* to be rankable if their labels (*y*_*i*_ and *y*_*j*_, respectively) satisfy

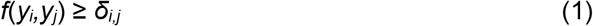

where *f* is a distance function and *δ*_*i,j*_ is the minimum necessary threshold of label separation. In classification, *f* was set to be an indicator function returning 0 if the arguments are identical and 1 otherwise, while *δ*_*i,j*_ was set to 0.5 for all (*i,j*). In linear regression, *f* was the L1-norm distance |*y*_*i*_ −*y*_*j*_| in the label space, and *δ*_*i,j*_ = max(*σ*_*i*_,*σ*_*j*_) was taken to be the expected measurement error, estimated by the standard deviations *σ*_*i*_ and *σ*_*j*_ computed across biological replicates.

Given the space of rankable pairs *R*, AUC is estimated by

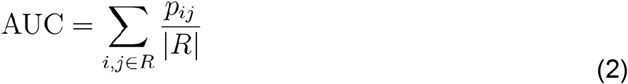

where *p*_*ij*_ is an indicator variable that takes on the value of 1 when the pair of samples *i,j* is correctly ranked by the predictor, and 0 otherwise.

### Outlier detection

We define the sample-specific AUC for the *k*-th sample as,

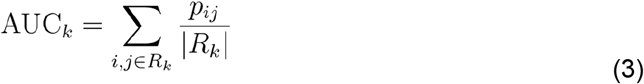

where *R*_*k*_ ⊂ *R* is the subset of all rankable pairs that include the sample *k*, and *p*_*i,j*_ has the same interpretation as in equation (2). Samples with significantly lower AUC_*k*_ than the overall AUC were considered to be potential outliers and inspected in more detail to decide whether they warrant an exclusion from the study. As with method comparison (**Figure 1c**), statistical significance was assessed by constructing a two-by-two contingency table cataloguing whether a given pair of samples is in *R*_*k*_ and whether that pair was ranked correctly by the corresponding model. Fisher’s exact test was used to determine whether pairs in *R*_*k*_ were ranked correctly significantly more often than pairs not in *R*_*k*_.

### Robust evaluation of predictors in the presence of confounders

To measure the effect of known confounders on the estimate of model performance, we considered a subset of rankable pairs where the difference in the confounder values was minimal. For discrete confounding variables (*e*.*g*., breast cancer subtype), the values were matched exactly. For continuous variables, we selected a single rankable pair per sample, such that the difference between the two values of the confounder was minimized. A possible unexplored alternative was to consider all samples that fell within a certain predefined “match” window for a given index sample. In all cases, we refer to resulting subsets of rankable pairs as *confounder-matched*, and the remaining rankable pairs as *confounder- mismatched*.

If AUC estimated on confounder-matched pairs was significantly lower than its equivalent derived from confounder-mismatched pairs, then this was interpreted as a strong indication that the corresponding predictor has learned to recognize the confounder instead of the variable of interest. Statistical significance was again assessed with a Fisher’s exact test applied to a two-by-two contingency table cataloguing whether rankable pairs were more likely to be ranked correctly if they are confounder-matched or confounder-mismatched.

### Combinatorial complexity considerations

While it is rare that all 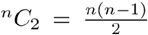 possible pairs of *n* samples will be rankable, the worst case complexity is nevertheless quadratic in *n* (**Figure S8a**). In small-sample studies, the O(*n*^2^) number of performance measurements produced by paired evaluation offers higher statistical power than the O(*n*) equivalent produced by traditional evaluation of applying a model to the entire test set at once. However, the quadratic complexity of paired evaluation is potentially prohibitive for evaluating predictive models on large-scale datasets, especially if LPOCV is employed. In such situations, the lack of statistical power is not usually a concern, and we propose to limit the total number of pairs considered in model evaluation by randomly selecting a single rankable pair for each sample (**Figure S8b**). A similar sampling procedure can also be used to define O(*n*) confounder-matched subsets (**Figure S8c**).

## Data availability

The results published here are in part based on data obtained from the AD Knowledge Portal (https://adknowledgeportal.org). We used the data available at https://github.com/labsyspharm/brca-profiling to train and evaluate predictors of drug sensitivity in breast cancer cell lines.

## Code availability

To employ paired evaluation in a LPOCV settings, we introduce a Python module lpocv that provides a flexible, automated way to identify sets of rankable pairs across a wide variety of machine learning settings and in the presence of known confounders. The rankable pairs can then be used to evaluate predictors, compare algorithms and identify outliers. The module is implemented following the object oriented programming paradigm and is compatible with algorithms in the scikit_learn package^38^, which is commonly used for machine learning tasks in Python. The implementation is open-source, with code publicly available on GitHub (https://github.com/labsyspharm/lpocv).

## Acknowledgments

We acknowledge support by the NIH Illuminating the Druggable Genome program (U24- DK116204), the NCI grant U54-CA225088 and the NIA grant R01 AG058063. ROSMAP study data were provided by the Rush Alzheimer’s Disease Center, Rush University Medical Center, Chicago.

## Supplementary information

**Figure S1:**
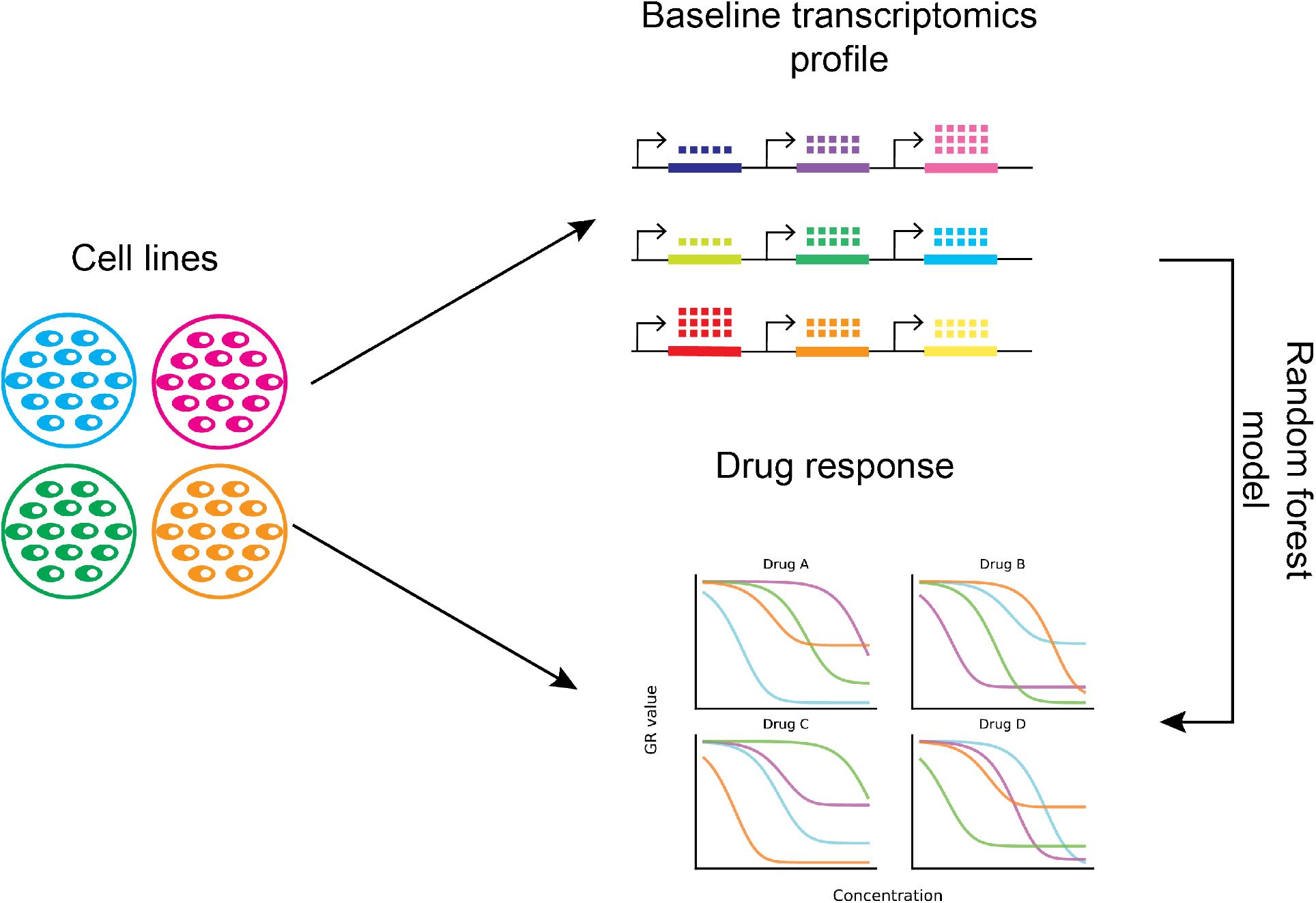
Random forest models are trained to predict the response of breast cancer cell lines to a panel of drugs. The response is measured as area over the growth rate curve. All models are given the expression of pre-selected genes as inputs. The expression is measured in baseline (i.e., pre-treatment) transcriptomic profiles.

**Figure S2:**
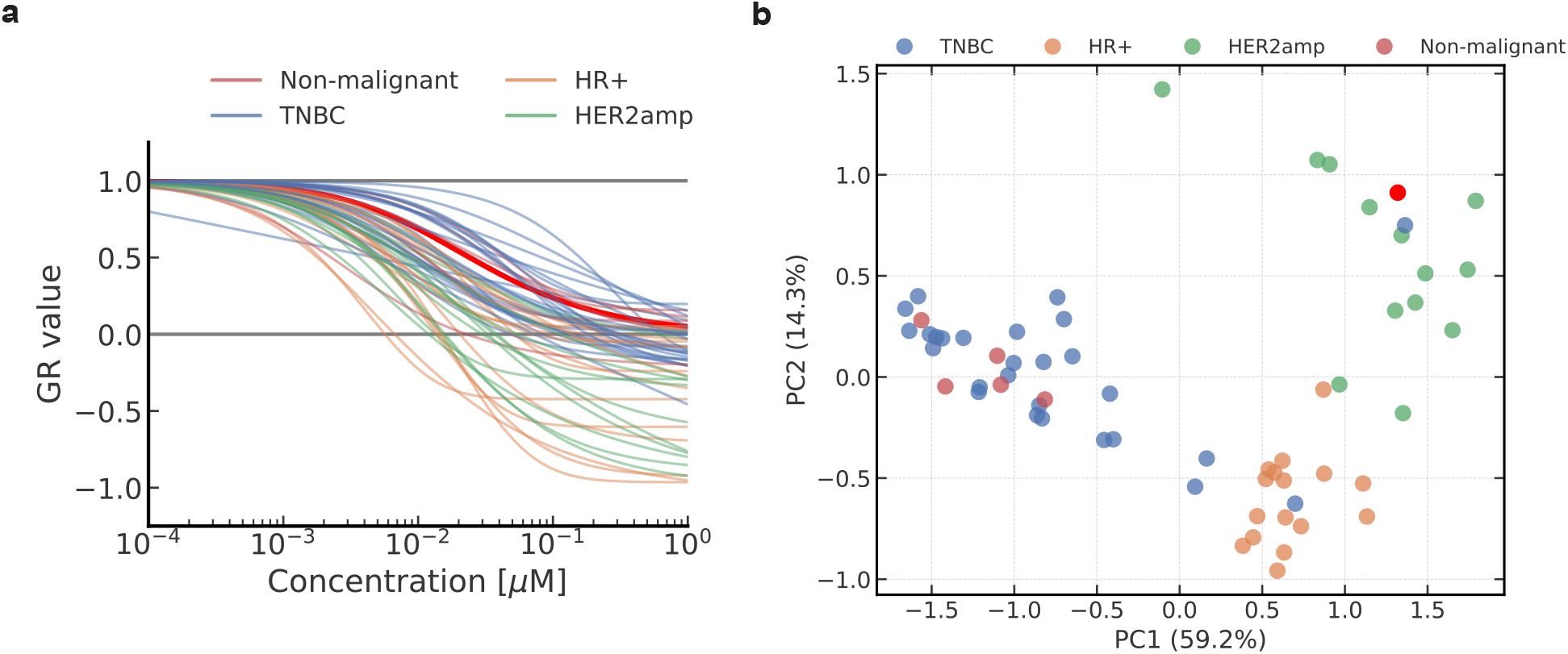
**a** Growth rates curves for torin2 across all breast cancer cell lines. **b** Principal components analysis of the baseline RNA-seq data, computed in the space of the top 20 most important genes (Figure 3). The “outlier” cell line ZR7530 is highlighted in red, all other cell lines are colored by their subtypes.

**Figure S3:**
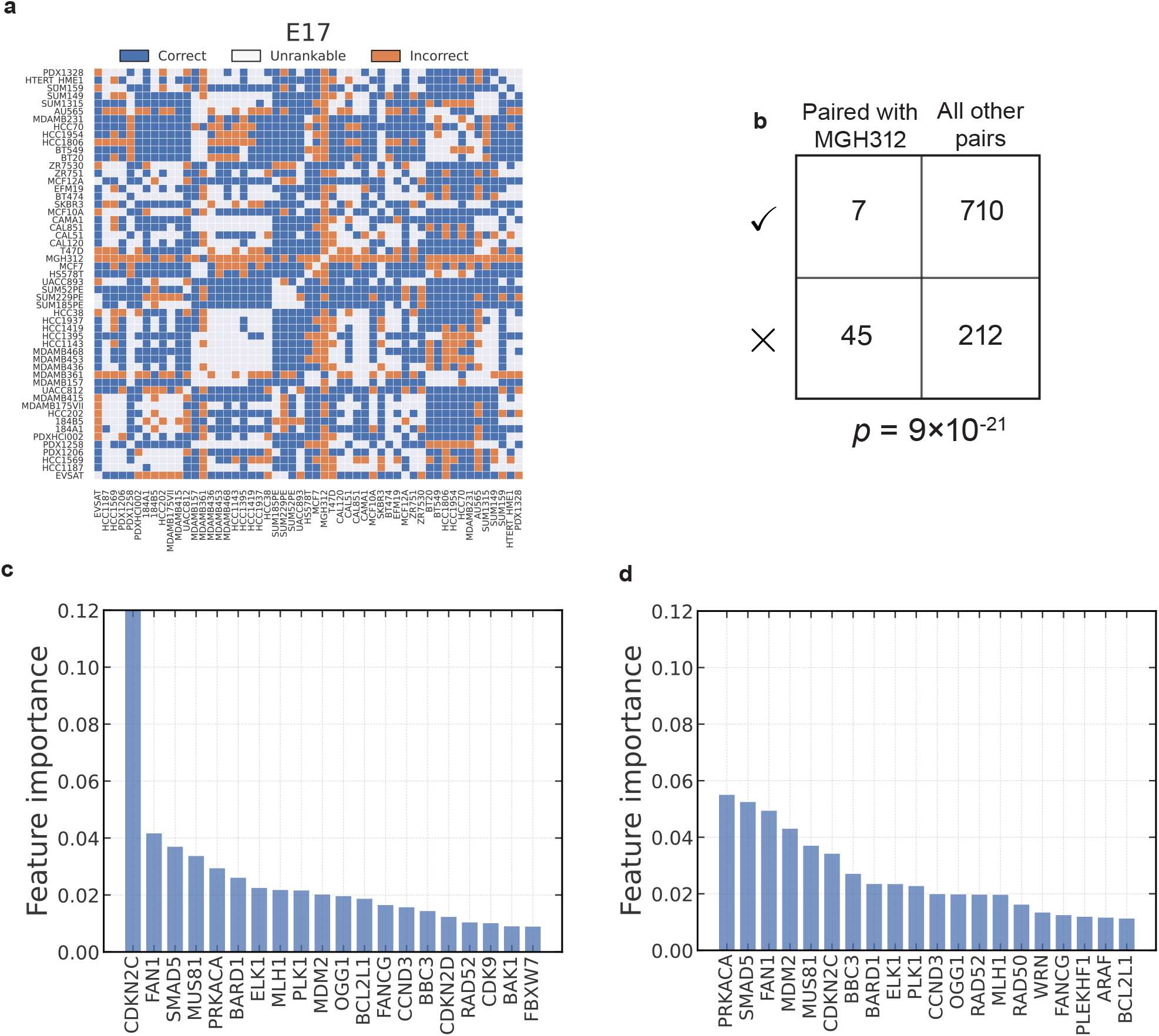
Outlier detection for E17. The interpretation of all panels is analogous to Figure 3.

**Figure S4:**
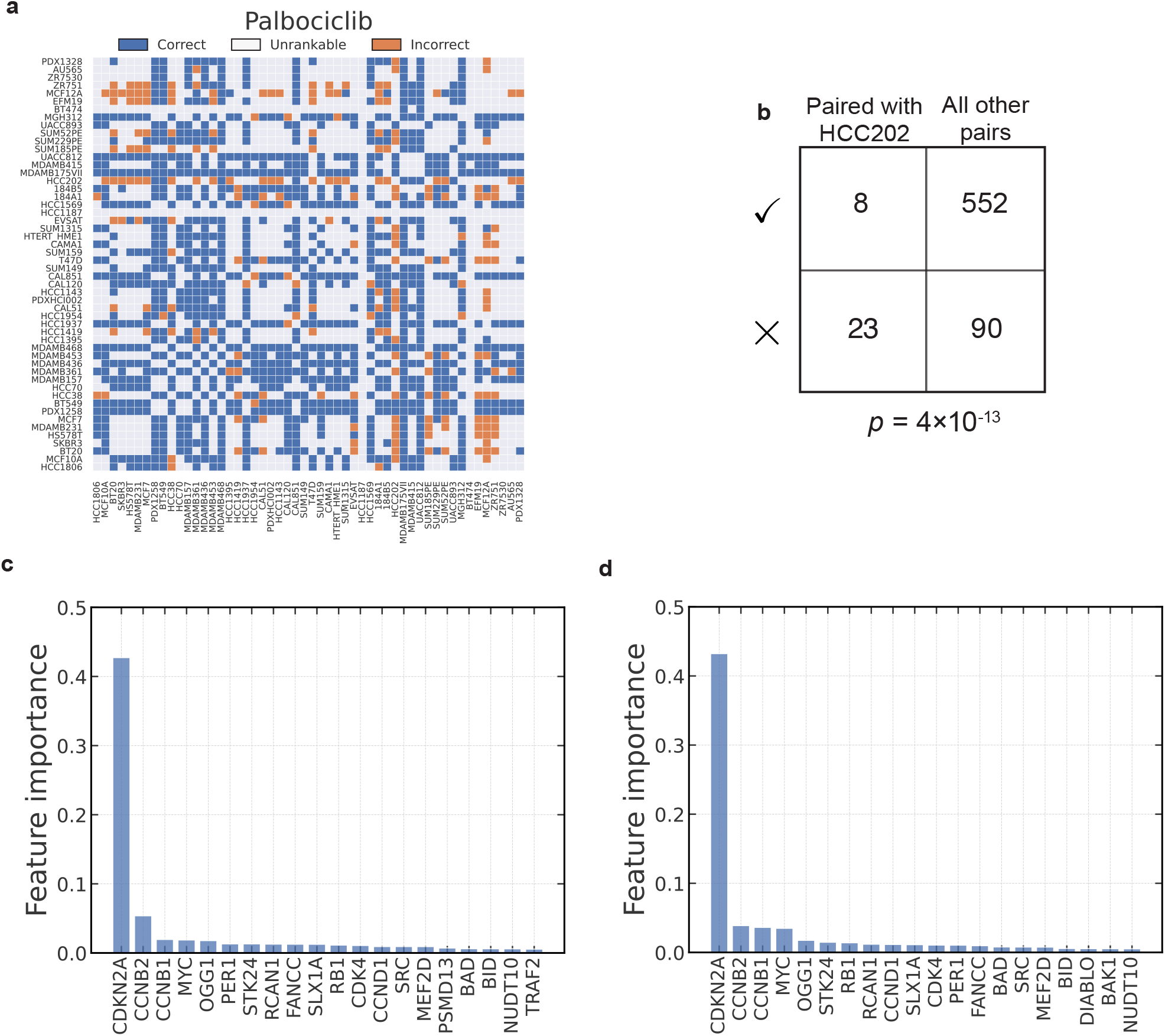
Outlier detection for palbociclib. The interpretation of all panels is analogous to Figure 3.

**Figure S5:**
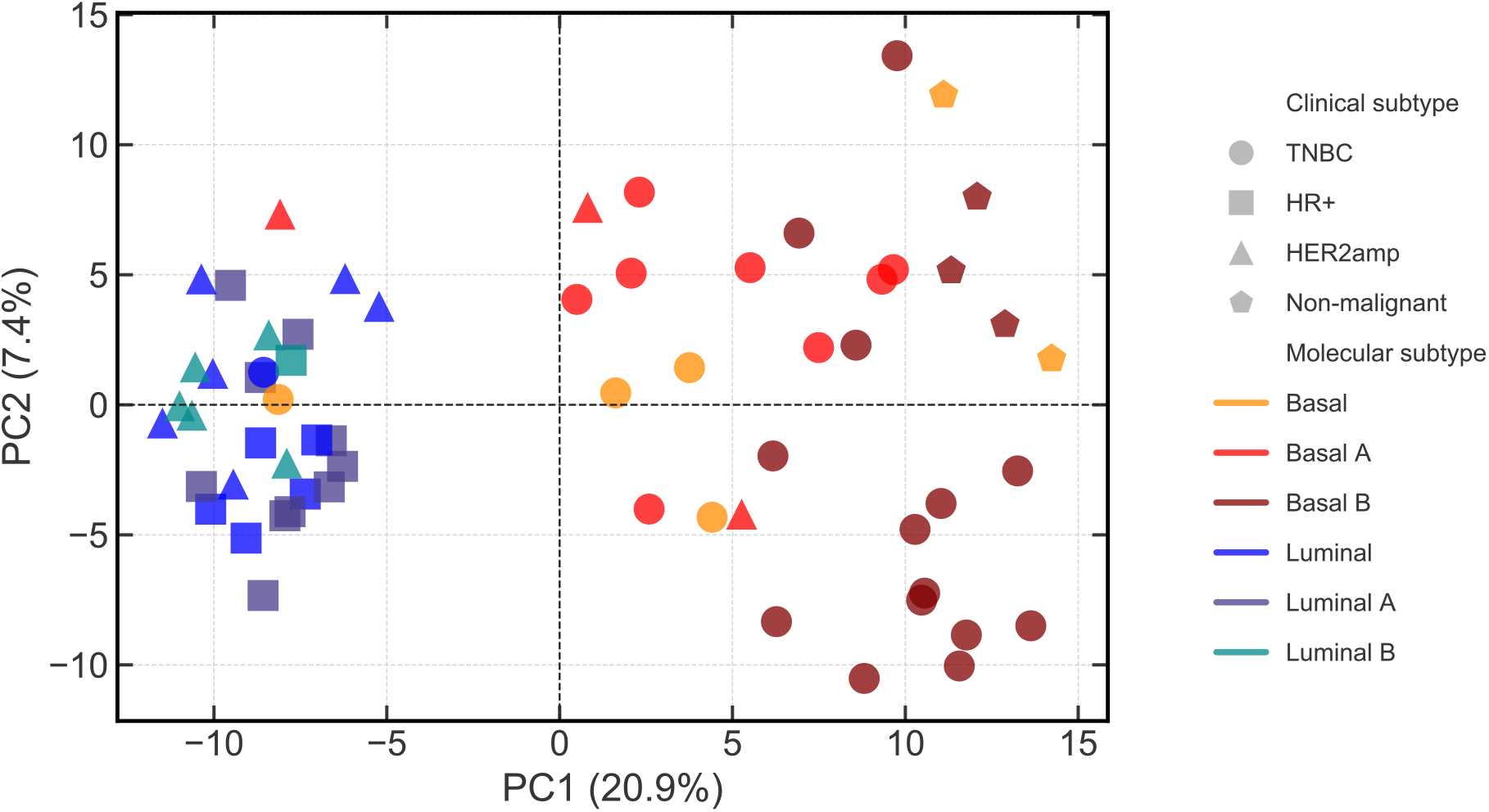
Principal components analysis of baseline RNAseq expression. The points represent individual breast cancer cell lines, shaped according to clinical subtype and colored by molecular subtype.

**Figure S6:**
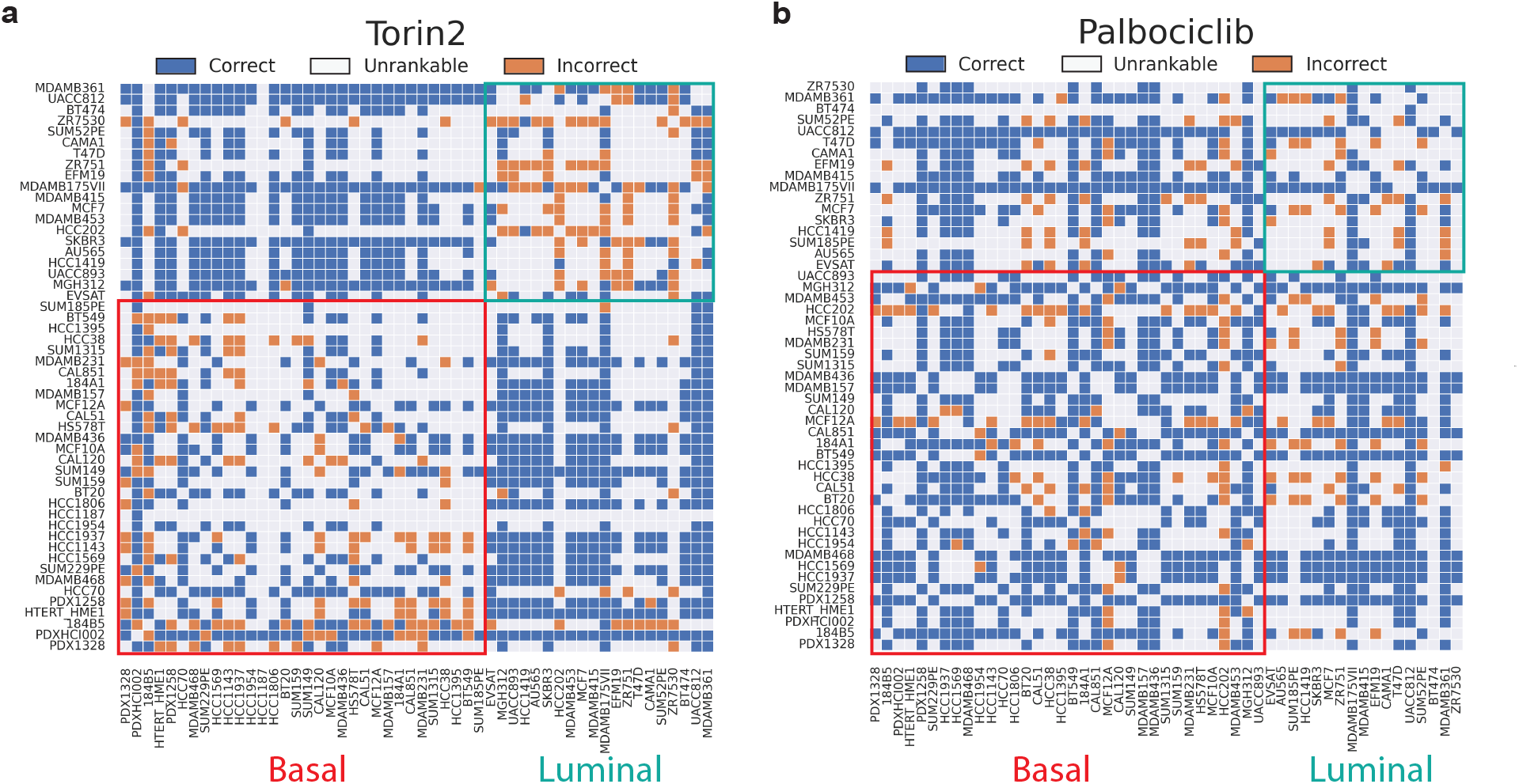
Effect of subtype in model prediction for torin2 and palbociclib.

**Figure S7:**
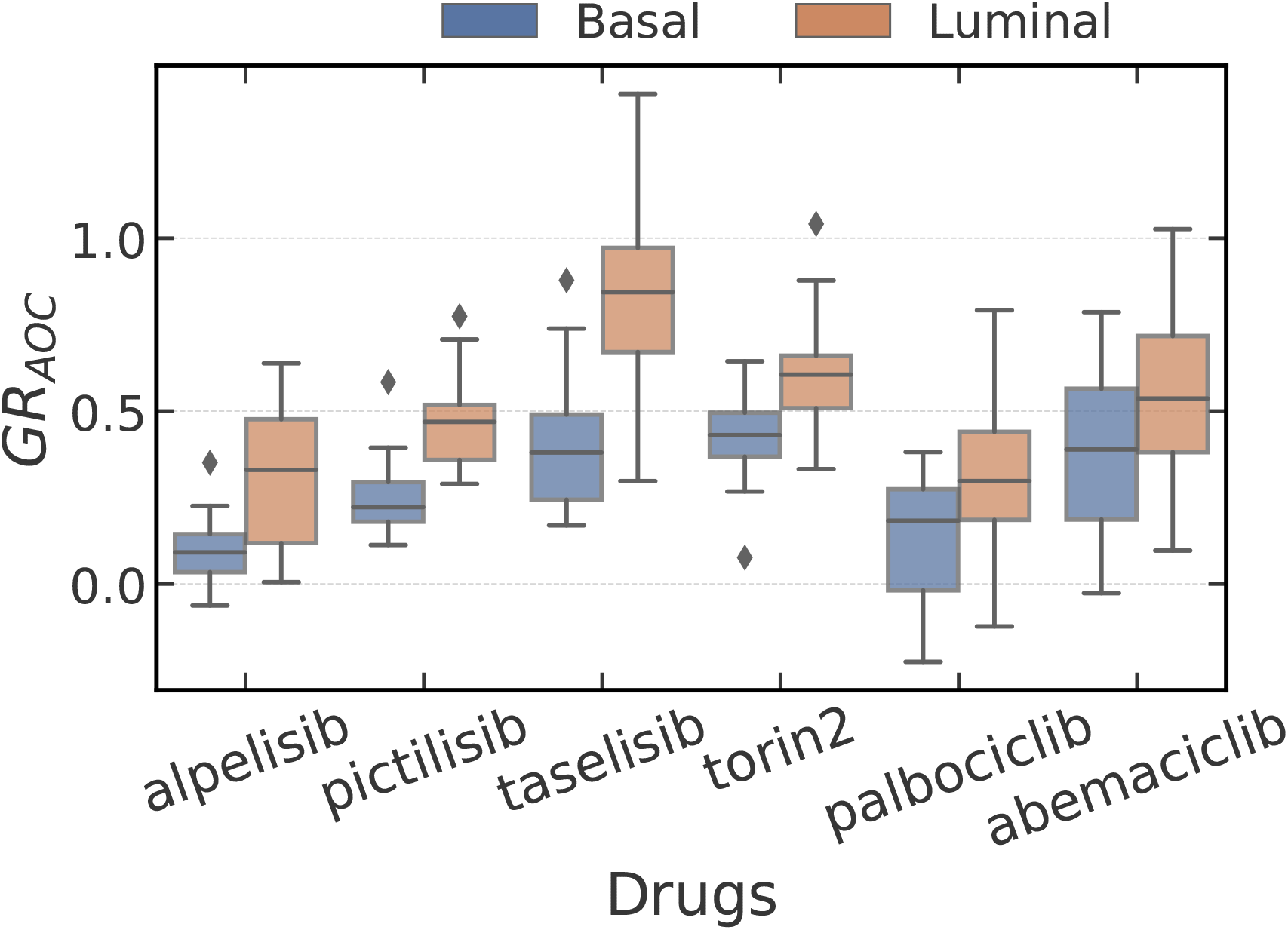
The distribution of *GR*_*AOC*_ for selected drugs. The values are plotted separately for basal (blue) and luminal (orange) cell lines.

**Figure S8:**
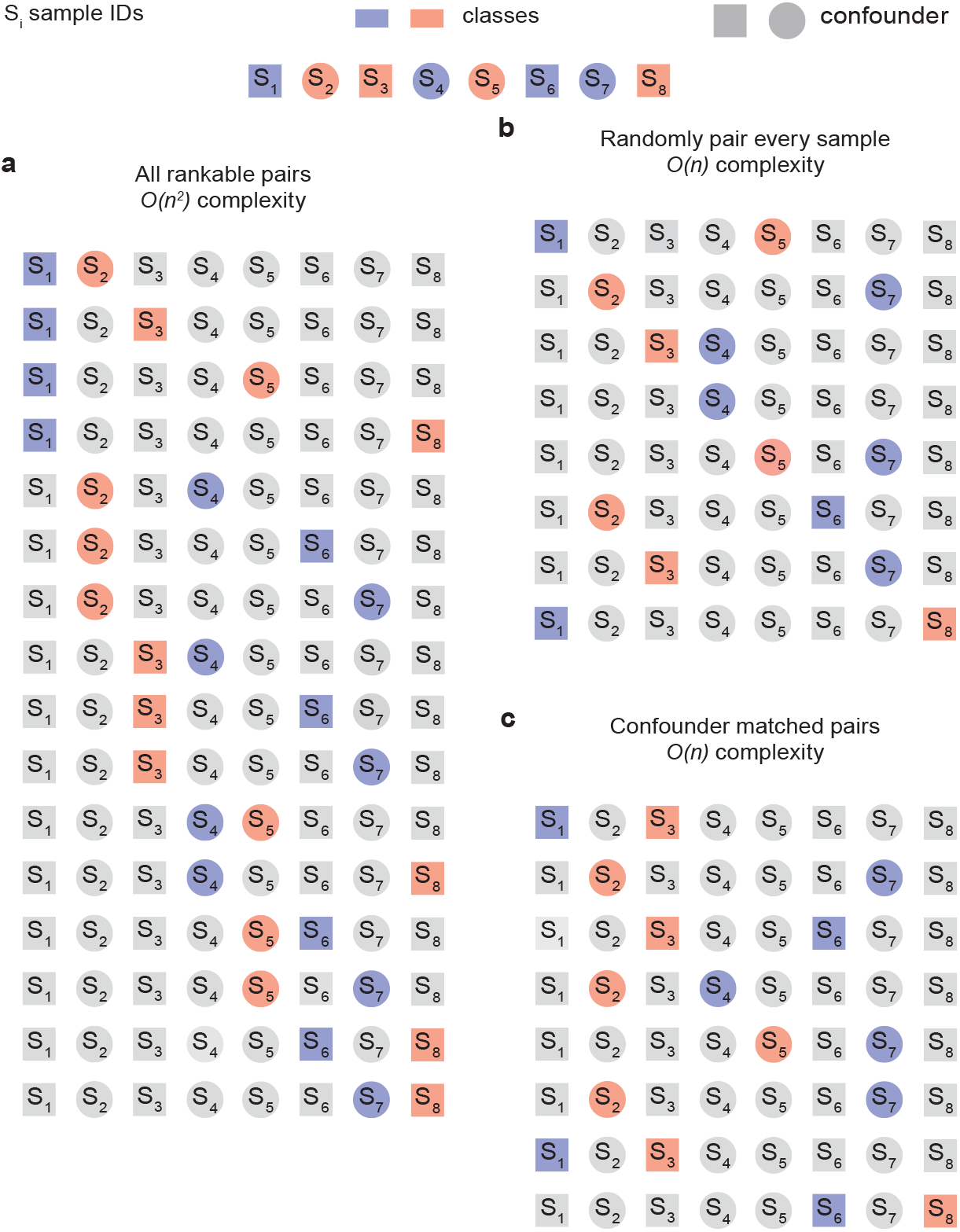
A schematic representation of how rankable pairs are defined in a dataset with *n* = 8 samples. Each square/circle element represents a sample. True labels are denoted with color, and the value of a binary confounder is represented by shapes. **a** All *𝒪* (*n*^2^) rankable pairs with a binary classification task. **b** An *𝒪* (*n*) subset of all rankable pairs, obtained by randomly selecting a single rankable pair for each sample. **c** An *𝒪* (*n*) subset of all rankable pairs, where each pair of samples is constrained to have the same value of the confounder.

**Supplementary Table 1:** The total number of rankable pairs used to evaluate predictors of drug sensitivity from mRNA expression.

| agent | num_pairs |
| --- | --- |
| Paclitaxel | 443 |
| Doxorubicin | 469 |
| Taselisib_GDC0032 | 714 |
| Pictilisib_GDC0941 | 358 |
| Torin2 | 389 |
| Vorinostat | 112 |
| Ipatasertib_GDC0068 | 333 |
| Everolimus | 535 |
| Tivantinib_ARQ197 | 125 |
| Cabozantinib | 66 |
| Saracatinib_AZD0530 | 296 |
| Dasatinib | 570 |
| Palbociclib_PD0332991 | 428 |
| Dinaciclib_SCH727965 | 224 |
| AZD7762 | 592 |
| Olaparib_AZD2281 | 50 |
| Alpelisib_BYL719 | 367 |
| A-1210477 | 7 |
| Buparlisib_NVP-BKM120 | 188 |
| INK128_MLN0128 | 526 |
| PF-4708671 | 65 |
| Neratinib_HKI272 | 588 |
| Cediranib_AZD2171 | 220 |
| Ceritinib_LDK378 | 205 |
| Trametinib_GSK1120212 | 460 |
| Luminespib_NVP-AUY922 | 322 |
| Abemaciclib_LY2835219 | 559 |
| Volasertib_BI6727 | 363 |
| ABT-737 | 131 |
| TGX221 | 190 |
| AZD1775 | 668 |
| AZD2014 | 693 |
| AZD5363 | 531 |
| AZD6738 | 294 |
| BJP-6-5-3 | 0 |
| BMS-265246 | 689 |
| BSJ-01-175 | 616 |
| BSJ-03-123 | 304 |
| BSJ-03-124 | 435 |
| BVD523 | 138 |
| CFI-400945 | 553 |
| E17 | 702 |
| FMF-03-145-1 | 705 |
| FMF-03-146-1 | 42 |
| FMF-04-107-2 | 787 |
| FMF-04-112-1 | 7 |
| Flavopiridol | 99 |
| GSK2334470 | 366 |
| LEE011_Ribociclib | 524 |
| LY2606368 | 747 |
| LY3023414 | 721 |
| MFH-2-90 | 837 |
| Pin1-3 | 50 |
| R0-3306 | 125 |
| Rucaparib | 198 |
| SHP099 | 61 |
| SY-1365 | 735 |
| THZ-P1-2 | 356 |
| THZ-P1-2R | 37 |
| THZ1 | 399 |
| THZ531 | 352 |
| YKL-5-124 | 570 |
| ZZ1-33B | 852 |
| senexin b | 150 |

